# Rewinding the molecular clock in the genus *Carabus* (Coleoptera: Carabidae) in light of fossil evidence and the Gondwana split: a re-analyses

**DOI:** 10.1101/2020.02.19.912543

**Authors:** Lars Opgenoorth, Sylvia Hofmann, Joachim Schmidt

## Abstract

1

**Background:** Molecular clocks have become powerful tools given increasing sequencing and fossil resources. However, outcome of calibration analyses depend on choosing priors. Here we revisit a seminal dating study of the genus *Carabus* by Andujar et al. proposing that their prior choices need re-evaluation with the hypothesis that reflecting fossil evidence and the Gondwanan split properly rewinds the molecular clock significantly. We used the same dataset including five mitochondrial and four nuclear DNA fragments with 7888 nt total length. We set the root age based on the fossil evidence of Harpalinae ground beetles in the Upper Cretaceous and introduce the Paleogene divergence of the outgroup taxa *Ceroglossus* (endemic to South-America) and *Pamborus* + *Maoripamborus* (Australia, New Zealand) as a new prior based on current paleontological and geological literature.

**Results:** The ultrametric time-calibrated tree of the extended nd5 dataset resulted in a median TMRCA *Carabus* age of 58.48 Ma (HPD95% 46.61-72.04), roughly 35 Ma older than in the Andujar study. The splits between *C. rugosus* and *C. morbillosus* (A), between *C. riffensis* from the European *Mesocarabus* (B), and between *Eurycarabus* and *Nesaeocarabus* (C) were dated to 19.19 (13.54-25.87), 25.95 (18.8-34.62), and 23.98 (17.28-31.47) Ma and were thus decidedly older than previously reported (7.48, 10.93, and 9.51 Ma). These changes were driven solely by constraining the Carabidae time tree root with Harpalinae amber fossils at ∼99 Ma. Utilizing the nd5 dating results of three well supported *Carabus* clades as secondary calibration points for the complete MIT-NUC data set lead to a TMRCA of *Carabus* of 53.56 (41.25-67.05) Ma compared to 25.16 (18.41-33.04) in Andujar’s study.

**Conclusion:** Taking into account the Gondwanan split as a new prior, together with the fossil evidence of the outgroup taxon Harpalini in the Late Cretaceous, our new approach supports an origin of the genus *Carabus* in the Paleocene-Early Eocene. Our results are preliminary due to the heavy reliance on the nd5 gene and thus will have to be tested with sufficient set of nuclear markers. In addition, uncertainties arise from dating the root age of the tree based on a single fossil and outgroup taxon which has a major effect on the results. Improvement of the fossil data base particularly in the supertribe Carabitae is thus strongly needed to reduce the currently large uncertainties in dating *Carabus* phylogeny.

## 2 Background

The molecular clock has become an increasingly powerful tool in biogeography and phylogenetics due to the ever-increasing genomic and fossil calibration data ^1^. However, phylogenetic dating is largely performed in Bayesian frameworks where the choice and number of calibration priors have a deciding impact on dating results ^1,2^. Consequently, there is often still a huge dating variance among studies even dealing with identical taxa and employing identical calibration points. Important factors for the disagreement are the placement of fossils in a given phylogeny and the handling of geological priors. The first is a matter of taxonomic discussion among species group specialists. Recent methodological improvements for better analyses of hidden characters in fossils like the usage of X-ray micro-computed tomography of amber inclusions to determine internal genital characteristics of tiny beetles ^3,4^ may help to resolve ambiguities in the long term. The handling of geological priors, on the other hand, is a broader discussion where the improvement could and should be somewhat more predictable and transparent across taxonomic groups. However, exactly in this part of the equation one can observe an almost arbitrary choice of geological sources and thus setting of respective molecular clocks ^5^. One classical geological event that has broadly left its imprint on biogeographic patterns is the split up and fragmentation of the Gondwanan landmasses ^6^. Studying the widely reviewed biogeographical literature dealing with the Gondwanan split it becomes evident that two general patterns emerge. Taxa that are good dispersers and occur on a broad range of terrestrial habitats have very diverse phylogeographic histories, often independent of the timing of the Gondwanan fragmentation. On the other side are taxa with poor dispersal capabilities and often very specific habitat preferences. Their evolutionary histories reflect the trademark vicariance pattern ^6^. Examples range from chironomid midges ^7^, stoneflies ^8^, scorpions ^9^, anurans ^10^ to plants such as *Nothofagus* ^11^, but see ^12^. Only for this second category (poor dispersers), the geological record is a means to calibrate the molecular clock.

Here, we want to revisit a seminal study for the calibration of the phylogeny of *Carabus* ground beetles ^13^ reflecting both, fossil evidence for the outgroup and recent geological as well as biogeographical consensus on the fragmentation of the Gondwanan landmasses. *Carabus* generally is described as a Holarctic genus that currently counts about 940 described species classified into 91 subgenera ^14^. Its diversification is bound to the Holarctic with a distribution throughout Eurasia, Japan, Iceland, the Canary Islands, North Africa, and North America ^14–18^. *Carabus* represents the most species diverse terminal clade of the “supertribe Carabitae” which also includes the Holarctic Cychrini, the Andean Ceroglossini (= *Ceroglossus*), the Australasian Pamborini (= *Pamborus* + *Maoripamborus*), and the cosmopolitan Calosomina (= *Calosoma sensu lato*). The latter was identified as the sister group of *Carabus* based on molecular data ^13,19–22^ which is in agreement with the morphological data ^15^.

Besides a Messinian fossil of *Carabus cancellatus* sensu lato ^23^, no additional fossil evidence is known for Carabitae older than the Pliocene and Quaternary periods. However, the poor fossil evidence certainly does not reflect the evolutionary age of the group. Based on an evolutionary model proposed by Terry L. Erwin in 1979 ^24^ the Carabitae represents a very old lineage of Geadephaga with its primary diversification reflecting continental drift events during the late Early Cretaceous. Penev et al. ^15^ propose that species belonging to the recent genera *Calosoma* and *Carabus* were present at least in the early Cenozoic time. The dates of the molecular phylogenetic study of Toussaint & Gillet^20^ correspond with these hypotheses, estimating the origin of Carabitae to about 170 Ma and the *Calosoma* - *Carabus* split to the Cretaceous. For divergence time estimation the split of Trachypachidae and Carabidae (estimated 200 Ma) was used from a reanalysis of the data of a previous study of McKenna et al. ^25^ using 34 Carabitae outgroup fossils ^26^. The most recent comprehensive analysis of Coleoptera molecular evolution was presented by McKenna et al. ^27^ and shows the Trachypachidae - Carabidae split at 170 Ma and the *Calosoma* - *Carabus* split in the Late Eocene and therewith, distinctly later as estimated by Toussaint & Gillet ^20^. For divergence time estimation McKenna et al. ^27^ selected 18 Carabitae outgroup fossils.

All these hypotheses are in more or less strong contrast to the molecular evolutionary models proposed by Andujar et al. ^13^ and Deuve et al. ^14^ with divergence time estimations mainly based on geological events. The latter authors propose the first diversification of Carabitae in the Paleocene-Eocene with a split of *Calosoma* and *Carabus* not until before the Oligocene. Such an Oligocene-Miocene emergence of the megadiverse genus *Carabus* is surprising with respect to the fossil evidence in the Carabidae family. Recent studies of Baltic amber inclusions make clear that representatives of modern ground beetle genera already existed during the Eocene, even those from the subfamily Harpalinae, with certain fossil species of the extant genera *Calathus* of the tribe Sphodrini ^28^, *Coptodera* of Lebiini ^29^, and *Limodromus* of Platynini ^30^. Also, the presence of Harpalinae is evident in the fossil record since the early Late Cretaceous ^31–33^. Finally, there is certain evidence from molecular genetic studies that Carabitae are phylogenetically older than Harpalinae ^34–36^, and that Harpalinae underwent rapid speciation in the Late Cretaceous and Early Cenozoic ^33,37,38^. In this regard, the question arises why the Carabitae would have undergone this markedly long phylogenetic standstill that would result from the timing proposed by Andujar et al. lasting a period of not less than 50-60 Ma.

This obvious dilemma leads us to revisit the dating background of the molecular study of the genus *Carabus* by Andujar et al. ^13^, not least because the evolutionary scenario proposed for this group was subsequently used by other researchers for dating approaches of their phylogenies of non-Carabitae taxa (e.g., ^39–42^). As we will show in the Material & Methods section in detail, Andujar et al. ^13^ did include several classical geological calibration events, namely the emergence of the Canary Islands, the Messinian salinity crisis i.e. opening of the strait of Gibraltar, and the disconnection of Japan from the mainland. In all three cases, we argue, that the approach chosen by them is not plausible from paleogeographical and paleoecological standpoints, respectively, but reflects a very common oversimplification of historical dispersal mechanisms. Also, we focus on another important issue of the Carabitae evolution: From a biogeographical point of view, particularly remarkable are the South American genus *Ceroglossus* and the Australasian genera *Pamborus* and *Maoripamborus*, which together form the sister clade to *Carabus* and Calosomina based on molecular data ^19,20^. A previous morphology-based hypothesis of the close relationship of *Pamborus* and *Maoripamborus* with the Cychrini tribe was identified to be a result of convergence ^43^. This view is also confirmed by the molecular data since Cychrini take the basal position within the “supertribe Carabitae” in previous studies ^13,20,34^. The split of the South American and Australasian taxa offers an additional possibility to calibrate the *Carabus* phylogeny. Since Andujar et al. ^13^ neglected this calibration point, we propose that their time tree massively underestimates the true age of the genus *Carabus* as was already presumed by Toussaint & Gillet ^20^.

In the light of these considerations, we hypothesize that i) adding a root age based on fossil evidence for Harpalinae and the inclusion of the Gondwanan split will push the dating of the crown age of *Carabus* to at least the Eocene and ii) a proper adjustment of the geological calibration points used by Andujar et al. ^13^ will resolve putative contradictions between those and the fossil data as well as the Gondwanan split. To test these hypotheses we here reanalyzed the datasets of Andujar et al. ^13^. Specifically, our new calibration strategy was based on a review of recent geological and biogeographical literature dealing with i) taxa that have a Gondwanan distribution, extracting minimum and maximum calibration ages; ii) the onset of the Canary hotspot using the taxonomic split between mainland and island taxa instead of island taxa only, and iii) the uplift of the North-African mountains. Finally, we will use our findings of the *Carabus* time tree to discuss the general need for a more differentiated and transparent usage of geological calibration points depending on life-history traits and habitat requirements of the taxa under study.

## 3 Methods

### 3.1 Phylogenetic data sets

For direct comparability of dating results, three phylogenetic data sets presented by Andujar ^13^ were utilized here based on the alignments deposited from Andujar and coworkers at treebase under the submission number 12410: http://purl.org/phylo/tree-base/phylows/study/TB2:S12410.

In short, the first data set is based on nd5 sequences from 58 Carabidae species and will be reffered to as the extended nd5 dataset. It includes 57 species of the supertribe Carabitae, with 51 species of the genus *Carabus* representing 16 subgenera and seven of the 13 main *Carabus* clades identified by Deuve et al. [14], one species each of the genera *Calosoma, Ceroglossus, Maoripamborus, Cychrus*, and two species of *Pamborus*, and the Harpalinae species *Abax parallelepipedus* as outgroup taxon.

The second data set included 34 specimens of the Carabidae family, including 19 species of the genus *Carabus*, two species of its sister taxon *Calosoma*, one species each of *Ceroglossus, Cychrus* (representatives of the supertribe Carabitae), as well as the Nebriitae species *Leistus spinibarbis* and the Harpalinae species *Laemostenus terricola* as outgroup taxa. Alignments were available for the mitochondrial genes *cox1-A, cox1-B, nd5, cytb, rrnL* and the nuclear genes *LSU-A, LSU-B, ITS2*, and *HUWE1*. A third data set is a subset of this second data set and only includes the ingroup, i.e. *Carabus* species.

### 3.2 Calibration strategy

In Table 2, the calibration scheme from Andujar et al. ^13^ used on the nd5 extended data set is shown. Table 3 shows the equivalent calibration scheme used in this study. Specifically, we follow the Andujar approach only in relation to the *Carabus cancellatus* fossil from France (F). Our treatment of the other calibration priors will be explained in the next paragraphs. Similar as in Andujar et al. ^13^ we used the resulting calibrated phylogeny to obtain ages for three well-supported cladogenetic events in the phylogeny of *Carabus* taking on the identical nomenclature for the respective splits between *Carabus* (*Macrothorax*) *rugosus* and *C*. (*Macrothorax*) *morbillosus* (Node A), *C.* (*Mesocarabus*) *riffensis* from the European *Mesocarabus* clade (Node B), and the split between the sister subgenera *Eurycarabus* and *Nesaeocarabus* (Node C). These nodes were selected by Andujar and coworkers because they are not (i) affected by systematic conflict, are (ii) old enough to avoid time dependence effects, and (iii) not excessively affected by saturation of molecular change. We used TreeStat 1.7.1 [48] to recover node ages from the sample of the MCMC search in BEAST and used the R MASS package ^44^ to obtain the gamma function with the “fitdistr” function (Table 4).

**Table 1.** Comparison of secondary calibration points derived in this study and in Andujar et al. 2011 for the calibration of the molecular phylogenies of single and combined datasets in *Carabus.* Median age and 95% HPD interval for the three clades were taken from the calibration analyses of the nd5 gene with BEAST2 (version 2.5.2) extracted with Tracer v 1.71. Gamma distribution was derived with the fitdistr function from the R package MASS. Four different sets of calibration points are shown in order of increasing age of the three clades: without the root and Gondwana prior, as in Andujar et. All, without the root calibration, and with all calibrations.

| NODE | CLADOGENETIC EVENT | Median Age and 95% HPD interval (Ma) |  |  |  |
| --- | --- | --- | --- | --- | --- |
|  |  | Without root and go calibration | Andujar et al. | Without root calibration | With all calibrations |
| A | Split between *C. (M.) rugosus* and *C. (M.) morbillosus* | 7.15<br>(3.02-13.49) | 7.48<br>(6.05-9.14) | 9.35<br>(6.73-12.50) | 20.276<br>(15.3-26.3) |
| B | Split of *C. (M.) riffensis* from European *Mesocarabus* | 9.61<br>(4.22-18.21) | 10.93<br>(8.90-13.26) | 12.81<br>(9.24-16.74) | 25.653<br>(19.8-32.7) |
| C | Split between *Eurycarabus* and *Nesaeocarabus* | 9.07<br>(3.64-16.71) | 9.51<br>(7.71-11.56) | 11.79<br>(8.73-15.51) | 23.276<br>(17.8-30.00) |

**Table 2.** Estimates of molecular age of the *Carabus* clade according to gene or gene set. Estimates were obtained from an MCMC run of 100 mio iterations (sampling parameter values and trees every 10,000 iterations, with a burn-in of 10%). With the exception of the evolutionary model used, which was inferred by bModelTest during the run, all parameters were the same for all analyses.

| gene (set) | This study |  |  | Andujar et al. 2012 |  |  |
| --- | --- | --- | --- | --- | --- | --- |
|  | Partition/<br>Clock | median / mean<br>age (Ma) | 95% HPD<br>interval (Ma) | Partition/<br>Clock | mean<br>age (Ma) | 95% HPD<br>interval (Ma) |
| *cox1-A* | 2P/ULN | 51.32 / 51.97 | 38.71-66.13 | 2P/SC | 19.79 | 15.2-24.9 |
| *cox1-B* | 2P/ULN | 49.62 / 50.2 | 37.54-64.15 | 2P/SC | 21.58 | 16.43-27.71 |
| *cytb* | 2P/ULN | 65.54 / 66.54 | 47.49-87.31 | 2P/SC | 25.77 | 19.74-32.91 |
| *nd5* | 2P/ULN | 53.07 / 53.67 | 39.95-68.14 | NP/SC | 20.71 | 151.9-26.15 |
| *rrNL* | 2P/ULN | 63.10 / 64.91 | 40.90-92.59 | NP/SC | 29.91 | 19.4-42.76 |
| *LSU-A* | NP/ULN | 35.01 / 38.44 | 20.29-65.48 | NP/ULN | 13.37 | 8.35-24.85 |
| *LSU-B* | NP/ULN | 59.44 / 64.15 | 30.56-108.45 | NP/ULN | 20.36 | 11.23-36.61 |
| *ITS2* | NP/ULN | 90.03 / 94.68 | 48.14-151.44 | NP/ULN | 31.17 | 16.8-52.47 |
| *HUWE1* | NP/ULN | 67.83 / 69.99 | 42.75-101.51 | NP/SC | 30.83 | 21.79-41.36 |
| *MIT* | 2P/ULN | 48.16 / 49.16 | 39.64-59.36 | G-2P/SC | 21.58 | 17.98-25.4 |
| *NUC* | NP/ULN | 72.02 / 74.85 | 43.25-112.86 | NP/ULN | 28.5 | 16.97-44.65 |
| *MIT NUC* | G-2P/ULN | 51.96 / 52.25 | 42.27-63.01 | G-2P/ULN | 25.16 | 18.41-33.04 |

**Table 3.** Calibration scheme from Andujar et al. 2012 ^13^. Denomination of the evolutionary events follows the Andujar study.

| Evolutionary event: node | Calibration event | Event age (Ma) | Priors on node ages | Prior 95% age interval |
| --- | --- | --- | --- | --- |
| **C** \| Split between two Canarian endemic species: *Carabus* (*Nesaeocarabus*) *coarctatus* and *C. (N.) abbreviatus* | Volcanic emergence of Gran Canaria | 14.5 | Uniform ($a = 0$, $b = 14.5$) | 0.03-14.14 |
| **F** \| *Carabus* (*Tachypus*) *cancellatus* fossil | Messinian deposits of Cantal (France) | 5 | Lognormal ($\mu = 25$, $\sigma = 1.5$, offset = 5) | 5.4-158.5 |
| **J1** \| Radiation of *Damaster* | Final disconnection of Japan from mainland | 3.5 | Truncated Normal ($\mu = 3.5$, $\sigma = 1$, $a = 0.1$, $b = 1000$) | 1.55-5.46 |
| **J2** \| Radiation of *Leptocarabus* | Final disconnection of Japan from mainland | 3.5 | Truncated Normal ($\mu = 3.5$, $\sigma = 1$, $a = 0.1$, $b = 1000$) | 1.55-5.46 |
| **J3** \| Radiation of *Ohomopterus* | Final disconnection of Japan from mainland | 3.5 | Truncated Normal ($\mu = 3.5$, $\sigma = 1$, $a = 0.1$, $b = 1000$) | 1.55-5.46 |
| **J4** \| Split between subgenus *Isiocarabus* and *Ohomopterus* | Initial disconnection of Japan from mainland | 15 | Normal ($\mu = 15$, $\sigma = 1$) | 13.04-16.96 |
| **M2** \| Split between *Carabus* (*Eurycarabus*) *genei* from Corsica and North African *Eurycarabus* | Opening Gibraltar strait | 5.33 | Exponential ($\mu = 0.5$, offset = 5.3) | 5.31-7.14 |
| Split between two *Carabus* (*Rhabdotocarabus*) *melancho-licus* subspecies: **M3** | Opening Gibraltar strait | 5.33 | Exponential ($\mu = 0.5$, offset = 5.3) | 5.31-7.14 |

**Table 4.** Calibration strategy of this study. Denomination of the evolutionary events follows the Andujar study ^13^ for better comparability.

| Evolutionary event: node | Calibration event | Event age (Ma) | Priors on node ages | Prior 95% age interval |
| --- | --- | --- | --- | --- |
| **C** \| Split between the Canarian endemic species *Carabus* (*Nesaeocarabus*) *coarctatus* and *C. (N.)* from the mainland *Eurycarabus*. | Volcanic emergence of the Canary Hotspot | 60 | Uniform (upper = 60, lower = 0, offset = 0) | 1.5 – 58.5 |
| **F** \| *Carabus* (*Tachypus*) *cancellatus* fossil | Messinian deposits of Cantal (France) | 5.3 | Lognormal ($\mu = 25$, $\sigma = 1.5$, offset = 5) | 5.43 – 159.0 |
| **M2** \| Split between *Carabus* (*Eurycarabus*) *genei* from Corsica and North African *Eurycarabus*. | Surface uplift of the North African mountain ranges | 15 | Lognormal ($\mu = 15$, $\sigma = 0.5$, offset = 0) | 4.97 – 35.30 |
| **M3** \| Split between two *Carabus* (*Rhabdotocarabus*) *melancholicus* subspecies | Surface uplift of the North African mountain ranges | 15 | Lognormal ($\mu = 15$, $\sigma = 0.5$, offset = 0) | 4.97 – 35.30 |
| **GO** \| Split from Pamborini and Ceroglossini | Split of Australia between Antarctica/South America | 32-50 | Lognormal ($\mu = 78$, $\sigma = 0.41$, offset=0) | 32.1 – 159.0 |
| **RO** \| Root | Fossil Oodini in Burmese amber | 99 | Lognormal ($\mu = 130$, $\sigma = 4.27$, offset = 98.17) | 98.2 – 159.0 |

#### 3.2.1 Canary Islands prior

Using endemic lineages from island hotspots to date phylogenetic trees remains a problematical approach because the true age of a lineage might be older than the islands themselves given their hotspot origin (see Heads 2014^45^ for a comprehensive discussion of this problem). In Andujar et al.^13^ the emergence of Gran Canaria (ca 14.5 Ma) was used to date the split of *Carabus* (*Nesaeocarabus*) *coarctatus* (Gran Canaria) and *C.* (*N*.) *abbreviates* (Tenerife). This approach is based on the hypothesis that divergence occurred immediately after the emergence of Gran Canaria and then only one of the species migrated to the later emerged Tenerife (11.9 Ma^46^) and at the same time went extinct in the original one. We propose that it is more parsimonious that the ancestral *Nesaeocarabus* species migrated to the Canary Hotspot before the emergence of Gran Canaria and Tenerife, respectively, and diversification occurred on one or more of the presumed older islands which are submerged today. Under this scenario, migration of *Nesaeocarabus* species to Gran Canaria and Tenerife was possible from any older island of which no date of their respective submergence is available. Consequently, using the emergence of only one of the recent islands to date splits within *Nesaeocarabus* is potentially misleading and needs to be relaxed to account for the alternative possibility. This is achieved by instead using the age of the Canary Hotspot and the split of Nesaeocarabus from the mainland Eurycarabus as island habitats could have been present since the upstart of the hotspot. Since the age of the Canary Hotspot has been proposed to be 60 Ma based on Kinematic studies ^47^ we set the prior on the split between Nesaeocarabus and Eurycarabus to 60 Ma using a uniform distribution as no additional information is available that would favour a specific time.

#### 3.2.2 Japan prior

We omit the four calibration points from Japan following the logic, that neither the final nor the initial disconnection of Japan from the mainland is mandatory for the radiation of the *Carabus* lineages *Damaster, Leptocarabus*, and *Ohomopterus*, and the split between *Isiocarabus* and *Ohomopterus*, respectively, but the availability of their habitat. The Asian Far East with the dense ensemble of the complex folding systems of Kamchatka, Sikhote Alin, the Sakhalin, and Korean peninsulas, the Japanese Islands, and the Kurile island arc, was geomorphologically highly diverse long before the initial disconnection of Japan ^48,49^. Therefore, during the Late Cenozoic, the occurrence of suitable habitats for temperate *Carabus* has to be assumed particularly along slopes of the many, more or less separated mountain arcs and volcanos of the area. Consequently, it is highly probable that separation of the lineages was linked to the particular geomorphology of the area and the resulting differences in the regional climate and therewith, has predated the splitting events of the Japanese Islands from continental Asia markedly. We, therefore, conclude that concerning the biogeographical history of ground beetles and other soil arthropods these geological events are not suitable calibration events. This conclusion is supported by the well-known fact that ground beetle faunas of the more or less directly adjacent mountain systems of Europe and continental Asia, which were not separated by sea during the Late Cenozoic, each are markedly differentiated. In this respect, the most impressive example is complex mountain ensemble of High Asia: The *Carabus* faunas of the Pamir, Greater Himalaya and Tibet (with the mountains of Western China) differ by nearly 100% even on the subgeneric level and is thus much more profoundly differentiated than that of the faunas of Japan and the Asian mainland ^50^. On the other hand, most of the main *Carabus* lineages were able to fly at least in the evolutionary history of the respective lineage, as evidenced by their phylogeny ^14^. For the early evolutionary history of the group, a sea can thus be considered a barrier with restricted efficacy for *Carabus* ground beetles. This becomes also obvious from the above-discussed occurrence of *Carabus* species on the Canary Islands because these islands never had a terrestrial connection to the continent.

#### 3.2.3 Gibraltar Strait vs. uplift of North African mountain ranges

The genus *Carabus* is an extratropical group of beetles with its representatives adapted to the warm (meridional) to cold temperate or subarctic climates. The species are mesophilic or hygrophilic and are absent in deserts. Temperature and humidity preferences of the beetles have to be considered when using occurrences of land bridges in the past such as the closing of the Gibraltar Strait to hypothesize dispersal events. So far, there is no paleoecological indication (let alone evidence) that the climate in the depression of the Gibraltar Strait was suitable for *Carabus* during the terrestrial development of the area in the Messinian. Instead, it was hot and dry, with the occurrence of spacious salt marshes, while warm temperate conditions and occurrence of mesophilic forests developed along slopes of the mountain belts ^51,52^. Mountains with various suitable habitats had been uplifted on both sides of the Gibraltar Strait much earlier^53,54^. Also, based on the current molecular phylogenetic data it has to be assumed that the ancestors of both *Carabus* subgenera *Eurycarabus* and *Rhapdotocarabus* were capable to fly since fully developed hind wings are occasionally present in *Carabus granulatus* of the more derived Digitulati group ^14^. Consequently, multiple developments of winglessness in the evolution of *Carabus* have to be inferred. If active dispersal by flight was possible for ancestors of the taxa in question, mountain uplift in the western Mediterranean region could have been an important event for the evolution of *Rhapdotocarabus* and *Eurycarabus* species. Consequently, this scenario is completely independent of the opening and closing of the Gibraltar Strait. Therefore, we constrained the splits in these groups with the age of surface uplift of the current Rif and Maghrebian Mountains. In the Rif rapid exhumation is assumed during the Late Oligocene - Early Miocene (27-18 Ma; Monie et al. 1994^55,56^), and in the Maghrebian extensional deformation probably occurred at 25-16 Ma ^57^. A significant height of these mountain ranges was thus certainly achieved in the Mid Miocene. Also, uplift of the immediately adjacent Atlas Mountains has been attributed to Cenozoic thickening of the crust and Middle to Late Miocene thinning of the mantle lithosphere related to a shallow mantle plume ^58,59^. A significant part of the paleoelevations have been attributed to this latter mechanism, namely a third of the mean altitude of 1,500 m in the western High Atlas, and half of the mean 2,000 m in the central High Atlas ^58^. We, therefore, conclude, that sufficient heights for suitable *Carabus* habitats were developed south of the Gibraltar Strait at least 15 Ma. We use a lognormal distribution since the arrival of the respective ancestors and separate evolution of the North African lineages could not have started before the North African mountains had reached into significant heights with suitable habitats.

#### 3.2.4 Split of Australia from Antarctica/South America

The breakup of Gondwana has been reflected in a number of ways in biogeographic and phylogeographic reconstructions. For example, Upchurch ^60^ proposed four broadly different models namely Samafrica model, Africa-first model, Pan-Gondwana model, and trans-oceanic dispersal. However, here we are only interested in the split of Australia from Antarctica or Antarctica and South America to reflect the split between *Pamborus* with its Australian distribution and *Maoripamborus* with its New Zealand distribution (together forming the tribe Pamborini) from *Ceroglossus* (forming the monotypic tribe Ceroglossini) with its South-American distribution. Numerous studies have found reticulate histories along with the split of Australia between the Late Cretaceous and the Eocene when a shallow seaway between Australia and Antarctica likely was the last land passage between these continents ^10^. Lately, it was proposed that this final split was a diachronous sea-floor spreading that started in the West 93-87 Ma, progressing to central Great Australia 85-83 Ma, followed by separation in the western Bight at ∼65 Ma and finalizing in the Terre Adelie-Otway region ∼50 Ma (summarized in ^61^). The same authors discuss that there still is large uncertainty in connection with this break-up history and propose that the oldest confident interpretations of magnetic seafloor anomalies date to ∼45 Ma when Australia and Antarctica finally drifted apart ^61^. However, the South Tasman Rise was already submerged between 50 to 32 Ma as deep as 1000 m ^62^ making passage difficult for beetles. In summary, we again implement a conservative dating approach concerning *Carabus* by choosing a calibration age with a minimum split at 32 Ma. Also, we set the maximum age to 159 Ma following Andujar et al. ^13^ and Deuve et al. ^14^. This date is attributed to the oldest fossils which most definitely belong to the Carabidae, and are described from the Upper Jurassic of Kazakhstan ^31^. The much older fossil *Lithorabus incertus* from the Lower Jurassic of Kyrgyzstan was also described within the Carabidae family ^63^, but it is based on a very poor imprint of few parts of the exoskeleton which is why the systematic assignment is rather doubtful. We chose a lognormal distribution (see Table 2).

#### 3.2.5 Fossil evidence for Harpalinae

As Carabitae outgroup taxon, Andujar et al. ^13^ used *Abax parallelepipedus* for their nd5 data analyses. The genus *Abax* Bonelli is representative of the Pterostichini tribe of the ground beetle subfamily Harpalinae *sensu* Crowson ^64^ which includes by far the most species of Carabidae. Morphological and molecular genetic phylogenies consistently indicate the terminal position of Harpalinae within Carabidae ^34,36,65^. Harpalinae fossils are described from Upper Cretaceous deposits in South Kazakhstan, Beleutin formation, Turonian (93.5-89.0 Ma ^63^), and from the Burmese amber (ca. 99 Ma ^32,33^). Therefore, we set the prior for the Harpalinae to 99 Ma (see Table 3), again with a maximum age of 158.5 following the same logic as described in 3.2.4.

### 3.3 Calibration analyses with an assessment of the input of each calibration point

All phylogenetic analyses were conducted with BEAST2 (version 2.5.2, ^66^) and were run on the CIPRES Cyberinfrastructure for Phylogenetic Research ^67^. To assess the relative contribution of each calibration point of the respective priors, we ran seven analyses by removing one of the individual calibration points from Table 3 in each of the runs, and one run with all priors included. Individual calibration point impact was assessed using violin plots produced with the R package vioplot ^68^. Since the root prior proofed to have such a decisive input on the overall dating, a second violinplot was drawn that excluded the root calibration in all other stepwise omissions. In all approaches, we constrained *Calosoma* to be the sister clade to *Carabus*, and used *Abax* as outgroup since these relationships have been well established ^34,65^. Nucleotide substitution models were inferred during the MCMC analysis with bModelTest package ^69^ implemented in BEAST2. Otherwise, we followed the settings as in ^13^ and used a Yule process as a model of speciation, a strict molecular clock, and a random tree as a starting tree. Each run was performed with 100 million generations, sampling 10.000 trees and with a burn-in set to 10% of the samples. Convergence and stationary levels were verified with Tracer v1.7.1 ^70^. We annotated the tree information with TreeAnnotator v.2.5.2 and visualized it with FigTree v.1.4.2.

### 3.4 Calibration analyses of each marker and their combination

The analyses of time-calibrated phylogenies employing nodes A-C generally followed Andujar et al.^13^. In short, median age and 95% HPD interval for the three clades were taken from the calibration analyses of the nd5 gene with BEAST2 (version 2.5.2) extracted with Tracer v 1.71. These were 20.276 (15.3-26.3), 25.653 (19.8-32.7), and 23.276 (17.8-30.0) respectively. Gamma distribution was derived with the fitdistr function from the R package MASS (Shape 52.599; Scale 0.389, Shape 61.144; Scale 0.423, Shape 56.317; Scale 0.417 for the nodes A, B, C). We then calculated time-calibrated phylogenetic reconstructions of each gene as well as the combinations of i) all mitochondrial genes, ii) all nuclear genes, and iii) all mitochondrial and nuclear genes were based on a run of 100 million generations. Subsequent steps followed the same protocol as in 3.3 (see above). Mean, standard error, highest posterior density intervals (HPD 95%), and effective sample size of likelihood, evolutionary rates, and the TMRCA of *Carabus* were inspected using Tracer 1.7.1. Consensus trees were obtained in TreeAnnotator 2.5.2 [44] using the median age option. In all instances an uncorrelated lognormal (ULN) relaxed clock was employed. All mitochondrial data sets were analysed under a 2P codon partition scheme with site models and clock models unlinked.

## 4 Results

### 4.1 Calibration analyses with the extended nd5 data set

In our calibration analyses, we used the expanded *nd5* gene dataset of Andujar et al. ^13^ and as therein implemented a strict clock and 2P codon partitioning. The ultrametric time-calibrated phylogenetic tree is shown in Figure 1 with a median TMRCA *Carabus* age of 58.48 Ma (HPD 95% 46.61–72.04 Ma) which was markedly older than the age obtained in the Andujar study (Table 1). The resulting mean molecular evolution rate was 0.0073 (95% HPD 0.0056–0.0091) substitutions per site per million years per lineage (subs/s/Ma/l) compared to 0.0154 (95% HPD 0.0112–0.0198) in ^13^. Nodes A, B, and C (Figure 1) had median ages of 19.19 Ma (A), 25.95 Ma (B), and 23.98 Ma (C) (95% HPD 13.54-25.87, 18.8-34.62, and 17.28-31.47) and were thus also decidedly older than in ^13^ (7.48 Ma, 10.93 Ma, and 9.51 Ma).

**Figure 1.**
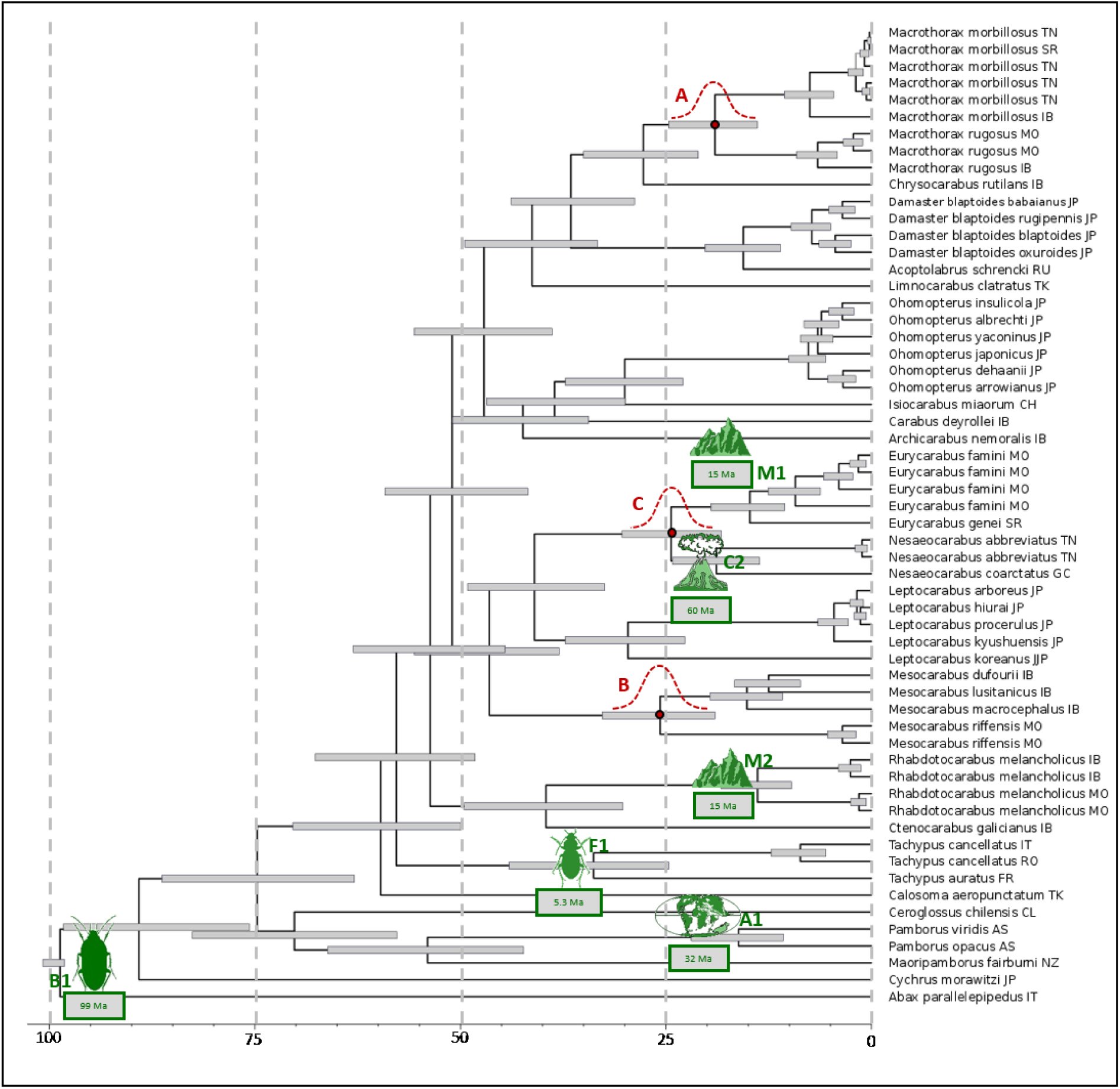
Ultametric time-calibrated phylogenetic tree obtained with BEAST2 for *nd5* data in *Carabus*. Green symbols depict calibration points employed – see **Table 3** for details. Red symbols depict cladogenetic events whose age distributions (see **Table 4**) have been utilized in subsequent analyses.

The violin plots in Figure 2 illustrate the overall impact of each calibration point on the timing of important phylogenetic lineages by depicting the age distribution when all points are used (all) and when each respective calibration point is left out of the calculation (-m2 through –root). As expected, most of this difference was attributed to the root prior which effectively overwrote any influence of other nodes as can be seen by the limited impact their omittance had on the dating results (Figure 2). This is further highlighted in Figure 3 which shows the violin plots entirely without the root prior. The graph also shows that the Gondwana calibration point had the next strongest impact on the dating, accounting for a median age shift for the Ceroglossini – Pamborini split of 10 My, thus mediating between the root and the remaining calibration points. More importantly, when included, clades A, B, and C are still ∼2 million years older than in Andujar (Table 1), while these clades were even younger than the Andujar results when the Gondwana prior was omitted in addition to the root prior.

**Figure 2.**
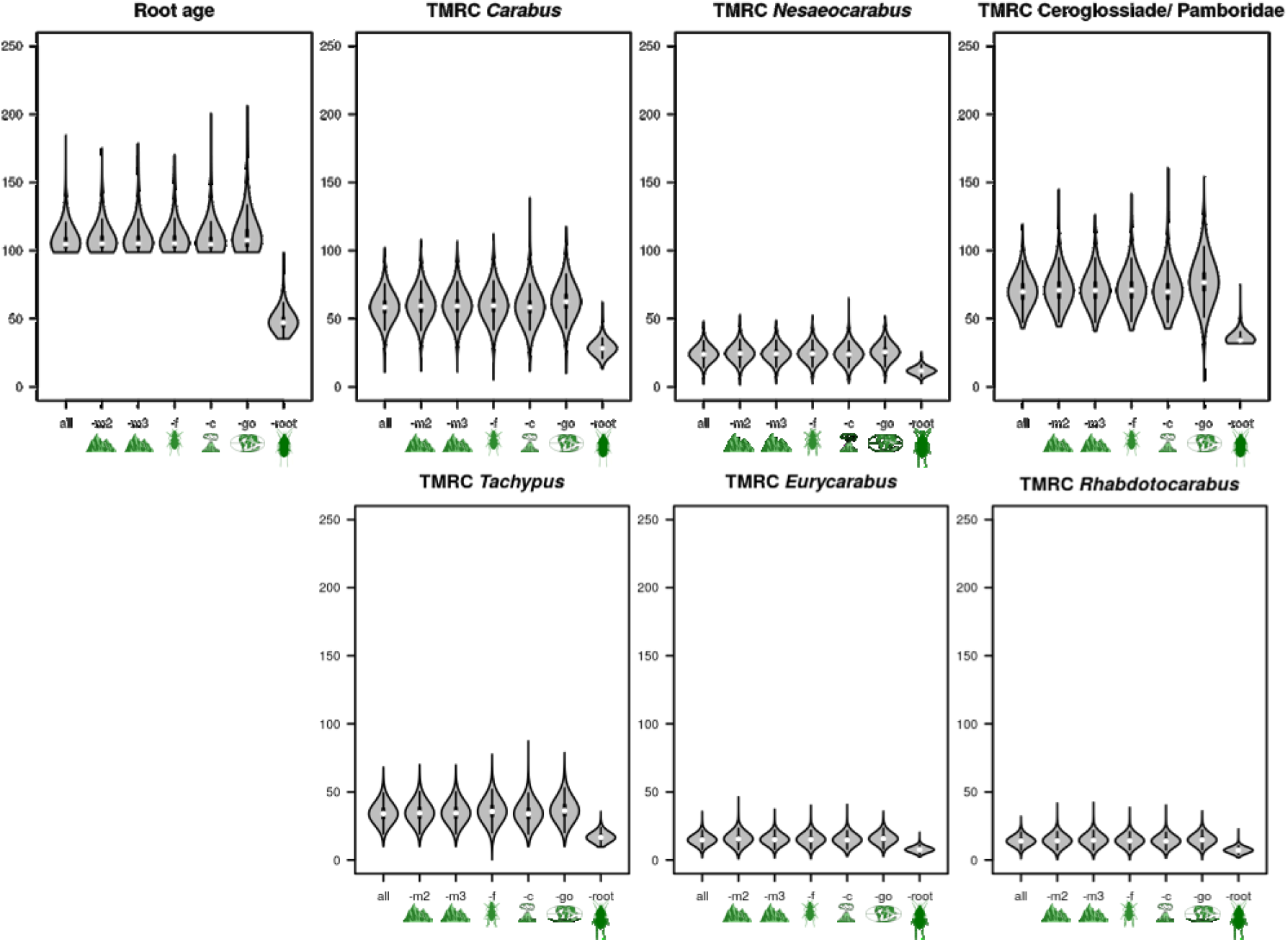
Violin plots of TMRCA ages in relation to individual calibration points. The plots depict mean (white dots), sd (black bars), 2*sd (black line), and density distribution (gray shape) of a clade’s age as obtained from the calibration analyses in **Figure 1** for each major clade in relation to exclusion (grey) and exclusive inclusion (orange) individual calibration points from the analysis. The x-axis denotes (letter) and depicts (icon) the respective calibration points in the respective analysis.

**Figure 3.**
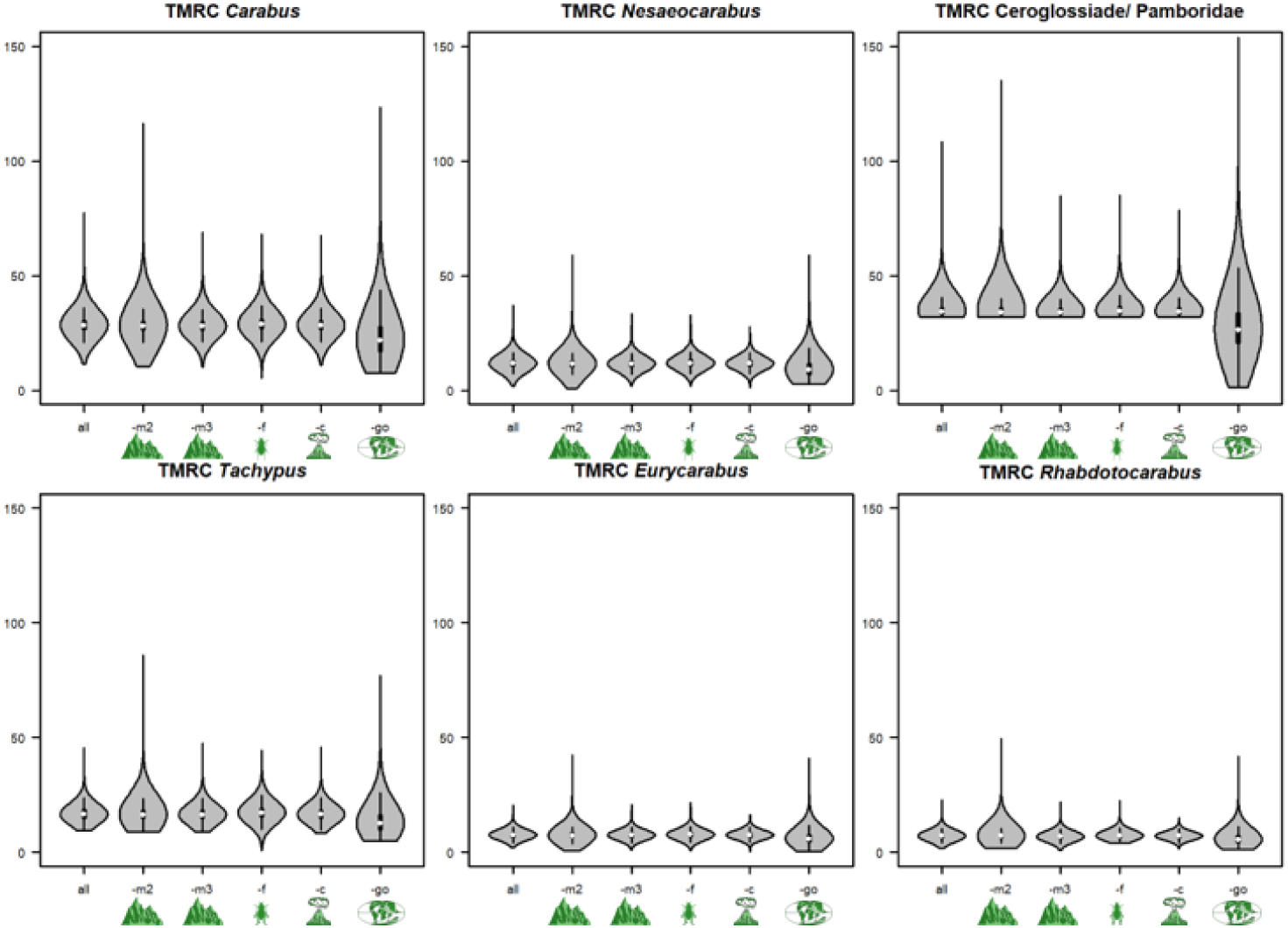
Violin plots of TMRCA ages in relation to individual calibration points when the root dating is omitted. The plots depict mean (white dots), sd (black bars), 2*sd (black line), and density distribution (gray shape) of a clade’s age as obtained from the calibration analyses in **Figure 1** for each major clade in relation to exclusion (grey) and exclusive inclusion (orange) individual calibration points from the analysis. The x-axis denotes (letter) and depicts (icon) the respective calibration points in the respective analysis.

As expected, the recovery of nodes generally had similar posterior probabilities (PP) as in the study of Andujar and coworkers thus, nodes A, B, C had PPs of 1.0 in all instances.

### 4.2 Time-calibrated gene trees and gene combination trees for the *Carabus* ingroup and for the Carabidae

Lineages A, B, and C were resolved in all gene trees and gene-combination trees except for LSU-A, and LSU-B where A-B, and B could not be resolved with adequate posterior probabilities, respectively. The derived median TMRCA of the *Carabus* clade varied markedly among genes ranging from 35.01 Ma for LSU-A to 90.03 Ma for ITS2. All medians and 95% HPD intervals for the respective genes and gene sets are given in Table 2. The ultrametric time-calibrated tree of the MIT-NUC data set lead to a TMRCA of *Carabus* of 53.56 (41.25-67.05) Ma compared to 25.16 (18.41-33.04) in Andujar’s study.

## 5 Discussion

### The time scale of *Carabus* evolution

We here reanalyzed the data presented by Andujar and coworkers^13^ in their study on the phylogenetic timing of the genus *Carabus* in the light of fossil evidence, the Gondwana split, and differing biogeographical interpretation of the geological record. We based our differning interpretation on a review of recent literature dealing with i) taxa that have a Gondwanan distribution, extracting minimum and maximum calibration ages; ii) the onset of the Canary hotspot, and iii) the uplift of the North African mountains. As expected, our study pushed the date of crown *Carabus* well beyond the Oligocene-Miocene dating of Andujar and coworkers to the Early Eocene when considering mitochondrial genes, and well into the Late Cretaceous when considering the nuclear genes of this study. Consequently, our findings support the model of Penev et al. ^15^ and Toussaint & Gillett ^26^ that species of *Carabus* were already present in the early Cenozoic. Also, our timing does not support evidence for a phylogenetic standstill of 50-60 My within Carabitae as it has to be assumed based on the data presented by Andujar et al.Figure 5. Furthermore, the Carabitae results now also fall in line with results from other taxa, for example, the dating of the Pelodryadinae-Phyllomedusinae split at 51.4 Ma (36.4-65.8) linked to the breakaway of Australia from South America ^10^. We consider this resemblance significant as Anura are known to show congruent evolutionary patterns with ground beetles based on similarly strong habitat ties ^71^ and thus this likely is not just a random match. However, we do have to stress that the rewinding effect is almost entirely based on the two fossils that were used to calibrate the root. Nevertheless, leaving these fossils out still led to a rewinding of the clock based on the inclusion of the Gondwanan split (Figure 3) though to a much lesser extend that would fail to reconcile the *Carabus* time tree with the existence of these fossils.

**Figure 5.**
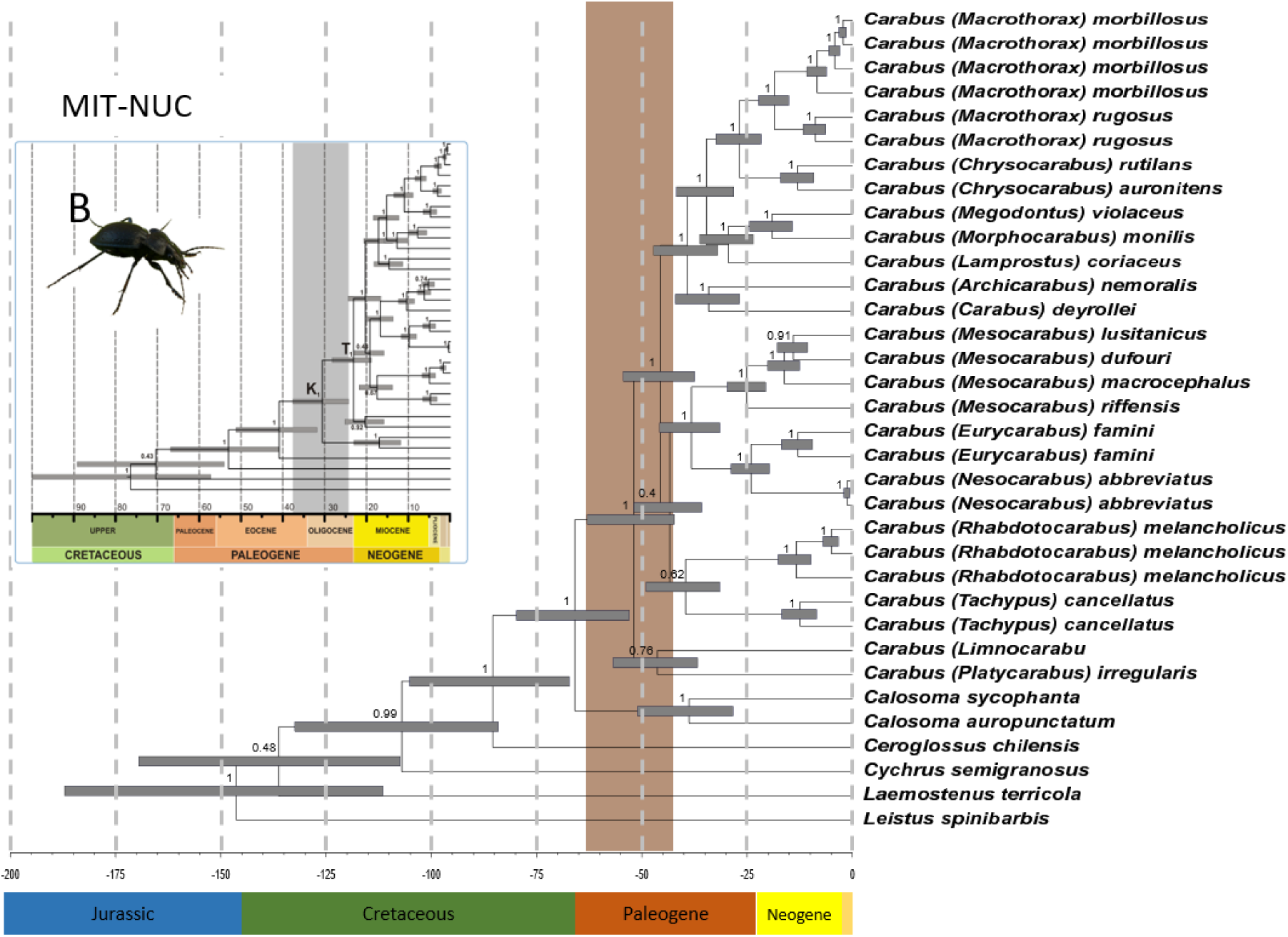
Ultrametric time-calibrated trees obtained with BEAST2 for each individual (left) and combined datasets (right) of the *Carabus* ingroup data set. Red clades and letters A, B, and C, are given for the individual gene trees when clades were supported by PP of more than 0.8, and for the latter likewise for the concatenated data sets and are aquivalent to the dated clades given in **Table 4**. Grey bars indicated the 95% HPD for the respective nodes, with the TMRCA bar being superposed for the respective phylogeny.

More specifically, when comparing the impact of the different added calibration points it is evident, that the inclusion of the Harpalinae fossils from Burmese amber as the root calibration has by far the strongest impact. Necessarily, this is true for the dating of the root itself, given that we implemented this fossil with a hard lower bound. In the extended nd5 calibration analyses this translates to a difference of almost 50 million years for the root when including or leaving this calibration point out (see Figure 2 and Table 2). Concerning the younger lineages, the root and the other calibration points produce more congruent results (Figure 2). The pattern has a similar but weaker trend when comparing the ultrametric time-calibrated trees obtained from the MIT data and the NUC or the combined MIT/NUC data set (Figure 4). This also implies that the dating of the internal clades will likely not be pushed back much further unless new older fossil evidence will appear. The initial split between *Carabus* and *Calosoma* is situated some 41 and 67 Ma (Andujar et al: 33-18 Ma ^13^) and thus occurred contemporarily with the final major phase of the breakup of Laurasia and opening of the northern Atlantic Ocean and therewith, with the split of the Nearctic and Palearctic regions in this part of the northern hemisphere. Andujar et al. ^13^ argued that the much younger *Carabus*/*Calosoma* split as derived from their analyses is congruent with the observation that *Carabus* is more diverse in the Palearctic region particularly due to low dispersal ability of the flightless species. However, the assumption of a flightless genus *Carabus* is misleading. Since functional hind wings are occasionally present in the extant species *Carabus* (*Limnocarabus*) *clatratus* and particularly in *C*. (*Carabus*) *granulatus* of the more terminal Digitulati group, flight capability must have been a trait not only for the most recent common ancestor (TMRCA) of the genus *Carabus* but for all TMRCA of its major lineages. Consequently, flight capability has to be assumed as an important precondition for the colonization of several marginal parts of the genus’ distributional area. This is particularly true in the south, such as the Canary Islands and the North African Mountains as discussed in the Material and Methods section of this study. Also it is very probably the precondition for the achievement of trans-Palearctic and trans-Holarctic distributions in several of the extant lineages (apart from those species which were dispersed by human activities just recently). The evolutionary events that originate the main extant lineages according to our data took place during the Mid and Late Paleogene and thus much earlier than estimated by Andujar et al. ^13^ and Deuve et al. ^14^. As such they are probably associated with the reorganization of the terrestrial biomes of the northern hemispheric regions due to climatic shifts ^72^ and major geomorphological events in Central and East Asia resulting from the uplift of the Himalaya-Tibet orogenic system ^73–76^. Since *Carabus* beetles are strictly adapted to the temperate climate and are thus absent in the tropics, climatic shifts might have had major impacts on the early distributional history of the genus, while the Neogene orogenetic evolution of the northern hemisphere was the main driver for allopatric diversification within the terminal lineages which resulted in an enormous number of wingless local endemic species particularly in the mountains of the mid-latitudes.

**Figure 4.**
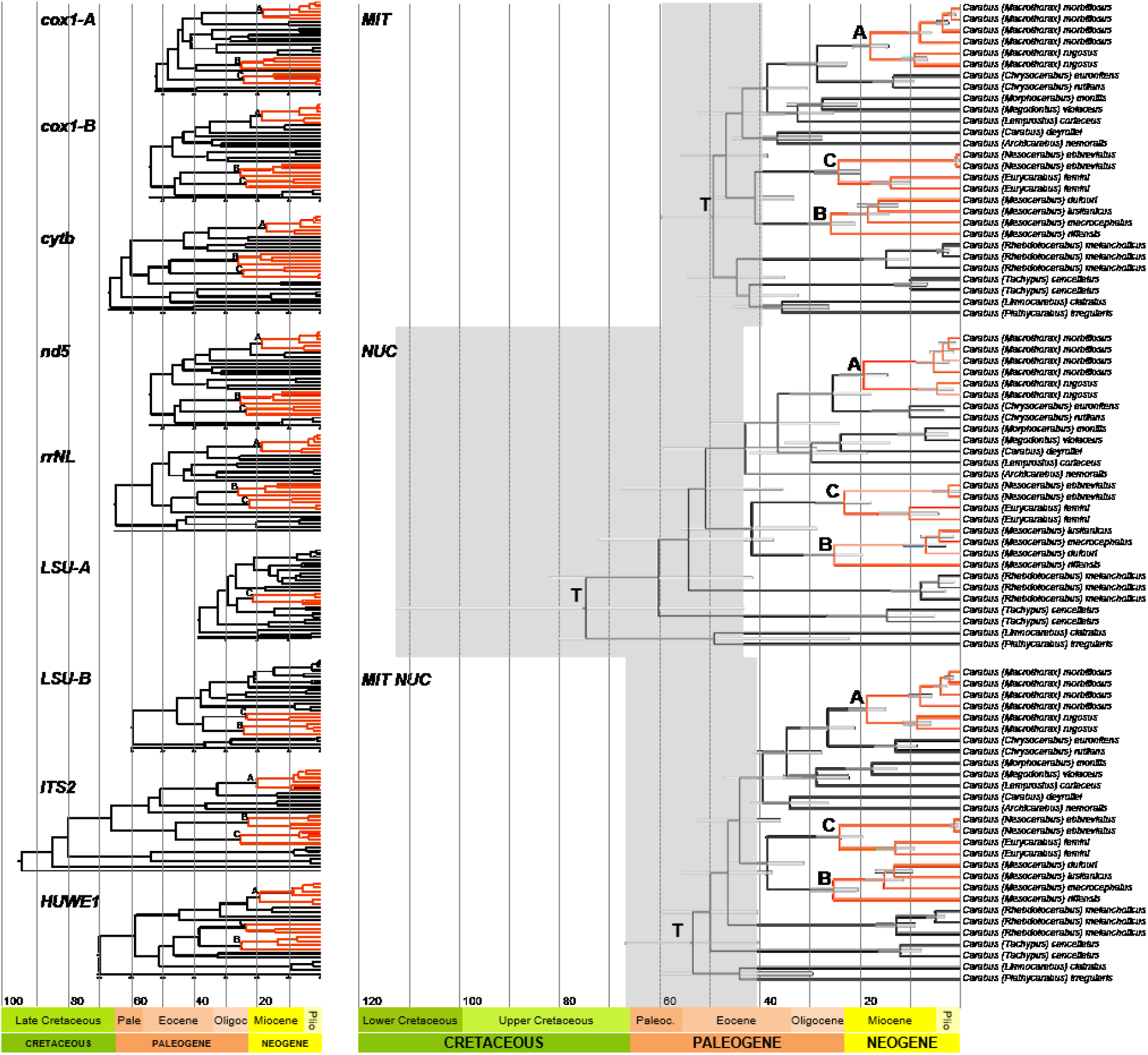
Ultrametric time-calibrated trees obtained with BEAST2 for each individual (left) and combined datasets (right) of the Carabus ingroup data set. Red clades and letters A, B, and C, are given for the individual gene trees when clades were supported by PP of more than 0.8, and for the latter likewise for the concatenated data sets and are equivalent to the dated clades given in Table 4. Grey bars indicated the 95% HPD for the respective nodes, with the TMRCA bar being superposed for the respective phylogeny.

Given the high impact of the root calibration on the dating results the future addition of Carabitae ingroup fossils originating from different geological periods to the analysis is mandatory. However, pre-Miocene Carabitae fossils are not found up to today. Concerning the markedly diverging *Carabus* – *Calosoma* divergence time estimations presented in the more comprehensive molecular phylogenetic beetle studies of Toussaint & Gillet ^20^: about 133 (98-172) Ma and McKenna et al. ^27^: 34 (18-57) Ma), which each are based on a large set of outgroup fossils, our results take an intermediate position and thus have some likelihood of standing up also in the future.

### Biogeographic Dating in the light of specific life history

Preferences for certain climatic conditions as well as dispersal ability of the species group have to be considered when geographical events are used to date phylogenetic events. Since *Carabus* is an extratropical genus with all its species being strictly adapted to the warm temperate or colder climates, there is no doubt about the origin of the genus *Carabus* in the northern parts of today’s Palearctic region. However, up to today, there is no clear evidence for a more detailed geographical origin. Previous molecular phylogenetic studies of the genus by Deuve et al. ^14^ show a simultaneous appearance of western (Arcifera, *Tachypus*) and eastern Palearctic elements (Crenolimbi, Spinulati) at the base of the *Carabus* tree. Reconstructing the early distributional history of the genus is more difficult since active dispersal by flight has to be assumed for the ancestors of most of the extant clades as discussed in the previous section. However, previous phylogenetic studies were probably biased by the author’s preoccupation with extant *Carabus* species being 99% wingless and thus underestimated the true historical dispersal ability of the species. E.g., the emergence of the subgenus *Nesaeocarabus* on the Canary Islands, which is one of the dating points in previous phylogenetic studies of the genus ^13,14^, has to be considered under the light of active dispersal from North Africa or southwestern Europe by flight. Dispersal by flight is likewise a most probable explanation for Europe-North Africa trans-Mediterranean distributional patterns as observed in the subgenera *Eurycarabus* and *Raptocarabus*. However, previous authors very probably assume dispersal “on foot” because of the desiccation of the Strait of Gibraltar during the Messinian crisis (about 6-5 Ma) and thus a short period of terrestrial connection of the continents was considered the prerequisite of the dispersal of the respective lineages ^13,14^. In the Material and Methods section, we have summarized arguments to reject this scenario since it is in strong contrast to the habitat preferences of the species. If active dispersal by flight is taken into account while discussing the dispersal history of the genus or using vicariance in biogeographic dating, the emergence and distribution of humid temperate habitats should be considered a much more important factor than terrestrial pathways. Consequently, concerning the trans-Mediterranean distributional patterns in some of the *Carabus* lineages, in pre-Pliocene times the distribution of mountainous areas as the provider of humid temperate habitats in the Mediterranean region, such as the North African mountain ranges, can be seen as the most important clue for the distribution of potential paleohabitats of the species.

## 6 Conclusions

Our study stresses the general need for a more differentiated and transparent usage of geological calibration points depending on life-history traits and habitat requirements of the taxa under study. Geological events need to be strictly interpreted from a biogeographic perspective including taxon-specific habitat suitability and dispersal abilities. Too often, geological events such as the Messinian salinity crisis or the opening/closing of the Strait of Gibraltar are transferred uncritically from a taxon where they might have enacted significant evolutionary pulses to taxa where they do not. Fossils play a crucial role as primary calibration points and much more effort should be invested in the future to improve the fossil database. In the here presented case of the dating of the *Carabus* clade, our study shows that the inclusion of fossil evidence in combination with taxon-specific biogeographic features results in a much earlier TMRC age of this clade. However, our conclusions should be considered preliminary due to the strong impact of the root calibration on the dating results. Additional Carabitae ingroup fossils are needed to prove our hypothesis, particularly those from pre-Miocene deposits.

## 7 Funding

The study was supported by Grants of the German Research Council (DFG) to Lars Opgenoorth (OP-219/2-1) and Joachim Schmidt (SCHM-3005/1-1).

### 7.1 Authors’ contributions

All three authors contributed to all parts of the manuscript and read and approved the final manuscript.

